# Enhancing spatial omics resolution by pseudo-interstitial pixels inference

**DOI:** 10.1101/2025.01.25.634874

**Authors:** Marco Antonio Mendoza-Parra, Maximilien Duvina, Ariel Galindo-Albarrán

## Abstract

**Motivation:** Spatially resolved omics technologies are enhancing our understanding of tissues architecture. Despite major technological improvements, gaining in spatial resolution becomes experimentally expensive, while generating spatial landscapes at moderate resolution combined with computational methods for depixelating data represent a cost-effective strategy allowing to enlarge the number of experiments to be performed.

**Results:** We have developed a computational strategy able to gain several-folds of resolution by inferring pseudo-interstitial pixels from their closest neighbors. This strategy has been validated in the context of public spatial transcriptomics data issued from melanoma, and human brain cortex tissue sections, by improving the identification of distinct tissue substructures. Furthermore, this methodology has been used for enhancing the resolution of consecutive sections collected from human brain organoids, as a way to demonstrate that a moderate resolution technology, combined with spatial depixelation processing allows to properly discern molecular tissue structures even in small tissues.

## 1 Introduction

Major developments in spatially-resolved transcriptomics (SrT) opened the way for the interrogation of the tissue architecture from the collection of molecular readouts across tissue sections. Recently, two orthogonal strategies emerged for addressing molecular tissue heterogeneity: (i) the use of unbiased strategies, based on the use of physical supports for either capturing the molecular readout with the help of positionally-indexed probes (Ståhl *et al*., 2016; Rodriques *et al*., 2019), or for transporting molecular barcodes across tissue sections like in the case of the use of microfluidics channels (Liu *et al*., 2020; Deng *et al*., 2022); (ii) the use of targeted strategies, based on the use of labelled probes for revealing spatial RNA location even at a subcellular level (Xia *et al*., 2019). Importantly, only the use of unbiased strategies provide means to discover new molecular targets, reason why major efforts in improving their resolution remains of interest.

While previous studies described integrative methods with single-cell transcriptomics data for performing cell-type heterogeneity deconvolution within spatial omics pixels (Longo *et al*., 2021), these approaches require ideally to count with single-cell data issued from the same tissue, implying performing further experimental assays.

Herein we introduce “MULTILAYER-expander”, a companion script to our previously described tool MULTILAYER (Moehlin, Mollet, *et al*., 2021; Moehlin, Koshy, *et al*., 2021), allowing to increase the spatial omics data resolution by inferring pseudo-interstitial pixels from their nearest neighbors. This strategy follows the concept behind image-depixelation methodologies used in large number of applications, including the up/down-scaling images for different screen resolutions, notably by Nearest Neighbor resampling (reviewed in (S Malini and Patil, 2018)). Furthermore, as part of the release of MULTILAYER-expander, we also provide an upgraded version of MULTILAYER, presenting an enhanced data display, an improved gene expression patterns detection, and a new module allowing to perform Uniform Manifold Approximation and Projection for Dimension Reduction (UMAP) and its projection within the spatial data.

## 2 Methods

### 2.1. Multilayer-expander implementation

Within spatial omics data, Multilayer-expander allows to create pseudo-interstitial pixels from their nearest neighbors. Specifically, Multilayer-expander proceeds as following for a newly inferred pseudo interstitial pixel to be located in position x&y:

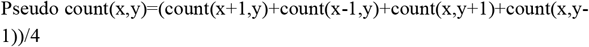

Where the inferred count is computed for each gene retrieved within the data. In this manner, inferred read counts per gene populate the new pixel. Importantly, Multilayer-expander performs this inference processing as many times as requested by the user, herein described as expansion steps. As input, Multilayer-expander requires a matrix containing physical coordinate identifiers as columns and gene IDs as rows. Users might also provide the number of required expansions (−n). Visium data requires to be converted to the aforementioned matrix format (a detailed procedure for this conversion is available within the documentation associated to Multilayer. Multilayer-expander has been developed in Python.

### 2.2. Multilayer.2025 version

Multilayer has been updated for providing a clearer display compatible with denser available data. Notably, the previously square-gridded view has been replaced by circular spots. In presence of interstitial data, Multilayer takes advantages of the new circular spots display for narrowing the physical space. Furthermore, Multilayer is now able to acquire for consecutive pixels even across interstitial maps. Finally, we have also included the possibility of generating dimensionality reduction UMAP displays within Multilayer (ScanPy package (Wolf *et al*., 2018)).

### 2.3. Data used in this study

Melanoma data has been obtained from the github repository of BayesSpace, notably because the original authors did not maintain their repository online: (http://www.ezstatconsulting.com/BayesSpace/articles/thrane_melanoma.html).

Dorsolateral prefrontal cortex data (slide n° 151673) has been obtained from the Leiber Institute (https://github.com/LieberInstitute/HumanPilot). Human brain organoids data were obtained from our previous publication (Lozachmeur *et al*., 2023). Furthermore, novel normalized spatial matrices as well as the corresponding depixelated processed data are available herein: Mendeley Data: https://data.mendeley.com/datasets/ptv86nczbv.

### 2.4. Dorsolateral prefrontal cortex data preprocessing

Data issued from the slide 151673 was converted from Visium 10x format to matrix format using the script “Visium-Converter” from Multilayer tool (GitHub - SysFate/MULTILAYER). The matrix was processed with the Multilayer-expander script by performing one expansion, and processed as following: For figures 3C&D, the matrix file was processed in R (R-project) using the “Seurat” package version 5.1 (Hao et al., 2024). First the matrix was converted to “MEX” format using the package “Matrix”, and read to generate the Seurat object. Second, the counts matrix was normalized by log normalization (NormalizeData), and the standard pipeline to generate the Uniform Manifold Approximation and Projection (UMAP). These steps were performed in a sequential way: FindVariableFeatures, ScaleData, RunPCA (npc=15), FindNeighbors (dims=1:15), RunUMAP, and the clusters were identified (KNN method) based on gene expression similarities using the command FindClusters at resolution “0.5”. These steps were applied in the matrix with or without expansion process, and graphics were generated with customized scripts in R language (pheatmap and geom_point from ggplot2). For figure 3G, the matrices with or without expansion were analyzed with Multilayer, as previously described in github repository: https://github.com/Sys-Fate/MULTILAYER. Selected genes were described in (Maynard *et al*., 2021).

### 2.5. Code availability

Both Multilayer-expander and the new version of Multilayer are available here: https://github.com/SysFate/MULTILAYER

## 3 Results

### 3.1 Enhanced tissue heterogeneity deconvolution by digital spatial omics depixelation processing

Inspired by methods used for image resolution scaling based on the Nearest Neighbor resampling algorithmic, we aimed at enhancing spatial omics resolution by inferring interstitial pixels from their closest neighbors. (**Figure 1A&B**). This is performed by assuming that cell heterogeneity is the consequence of gene expression complexity (Dries *et al*., 2021), which can be inferred in the newly interstitial pixels by computing gene expression average from the closest neighbor pixels. Importantly, this interstitial inference can be performed over multiple rounds of expansion (**Figure 1C**), and can be performed from spatial matrix data presenting linear pixels organization (e.g. Spatial Transcriptomics technology initially described by (Ståhl *et al*., 2016)), but also from physically interstitially organized data like those issued from the commercial platform Visium 10XGenomics.

**Figure 1.**
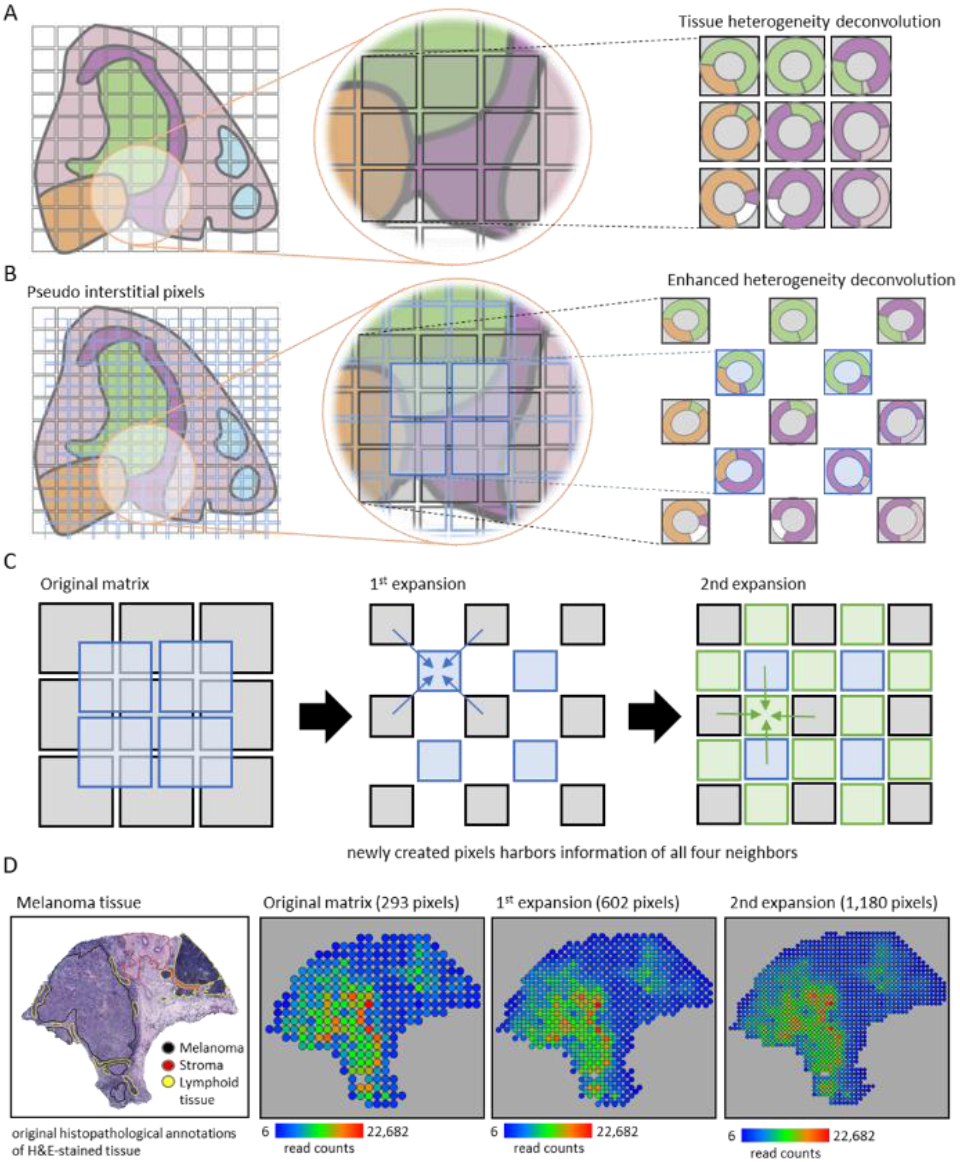
Enhancing spatial omics data resolution by inferring pseudo-interstitial pixels. **(A)** Scheme illustrating how a physical grid can capture the information associated to the tissue heterogeneity architecture. **(B)** Creating pseudo-interstitial pixels allows to further stratify the physical grid allowing to infer tissue heterogeneity information across consecutive pixels. Tissue scheme in (A) and (B) was adapted from (*Dries et al*., *2021*). **(C)** Newly created pixels are composed by the average read counts per gene assessed from the nearest neighbors, and are physically displayed within the source of information. In a first expansion process a linear matrix become interstitial and in a second round of expansion a larger linear matrix is retrieved. This process can also start from an interstitial matrix (e.g. Visium data). **(D)** Example of two rounds of pixel density expansion over a melanoma tissue data obtained from the public domain (*Thrane et al*., *2018*).

To validate the performance of this computational strategy we have analyzed a melanoma SrT data initially generated by (Thrane *et al*., 2018) with DNA arrays issued from the seminal technology developed by (Ståhl *et al*., 2016). This original data is composed by 293 pixels covering the melanoma tissue at a resolution of 100 microns (**Figure 1D**). By applying the spatial omics depixelation strategy across all genes in this map we have increased the pixel density to 602 in the first expansion process, then to 1,180 pixels after a second round of pseudo-interstitial pixels inference, which corresponds to a 4-fold gain in resolution. Importantly, this procedure did not distort the total read count maps, but as expected, it enhanced the visual rendering of the molecular tissue landscape (**Figure 1D**).

The histopathological manual annotation presented in the original article describing the presence of melanoma, stroma and lymphoid tissue (**Figure 1D**) was further characterized in a previous study using Bayesian statistics to achieve subspot level resolution (Zhao *et al*., 2021)). This methodology, named BayesSpace, allowed to enhance the identification of lymphoid tissue around the tumor border and potential immune infiltration, barely detected in the original data (Zhao *et al*., 2021). In agreement with these findings, our spatial omics depixelation strategy allows to visualize dense and continuous patches of pixels corresponding to over-expressing gene-markers related to chemokine activity (CXCL9, CXCL10) surrounding the tumor cell area (defined by PMEL over-expression) (**Figure 2**). Furthermore, regions associated with B cells (CD19 over-expression), Macrophages (CD14 over-expression) and even more clearly with T cells (CD3D over-expression) are better defined in the depixelated maps, demonstrating the advantage of increasing data resolution by incorporating pseudo-interstitial pixels.

**Figure 2.**
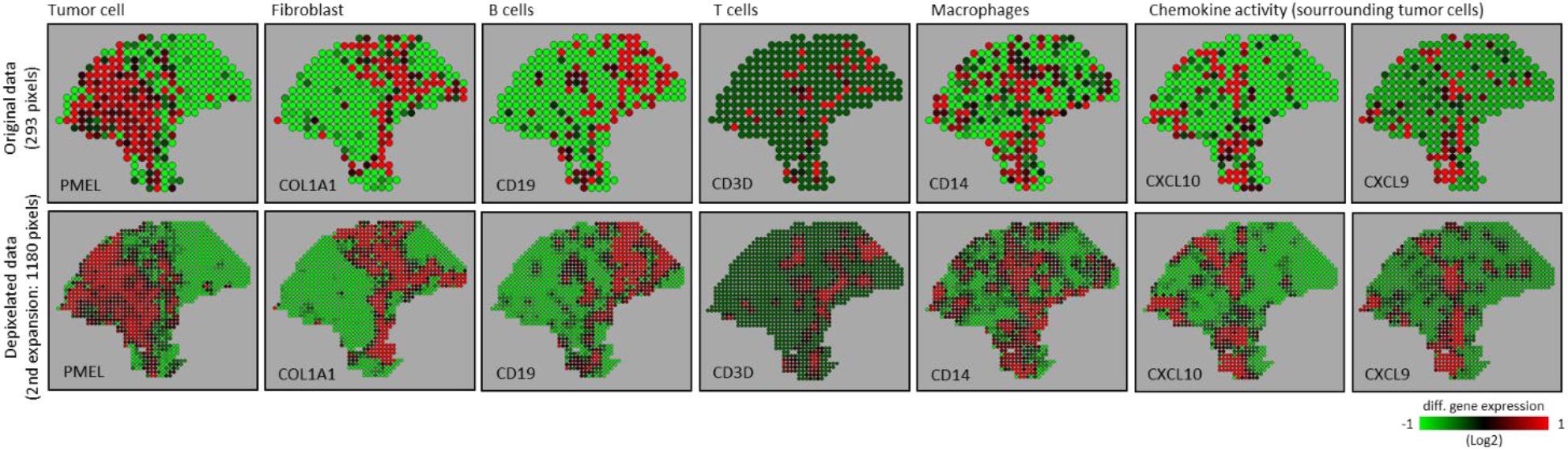
Enhanced cell-type signatures identification in melanoma tissue thanks to spatial depixelation processing. Gene marker over-expression associated to tumor cells, fibroblasts, B cells, T cells, Macrophages and chemokines are displayed. Differential gene expression displays were generated with Multilayer.2025.

### 3.2 Depixelation data processing enhances the identification of known cortical layers in human brain datasets

One of the main advantages of gaining in data resolution is the possibility of better resolve tissue substructures due to the fact of counting with larger number of pixels for data stratification. To address this point, we have used a dorsolateral prefrontal cortex tissue section data generated by (Maynard *et al*., 2021) with the Visium spatial expression profile technology (**Figure 3A**). This data has been used by previous studies to evaluate the performance of computational strategies to identify distinct layer-specific expression profiles, notably because this tissue presents six cortical layers and part of the white matter which were properly defined by manual annotation (**Figure 3B**).

**Figure 3.**
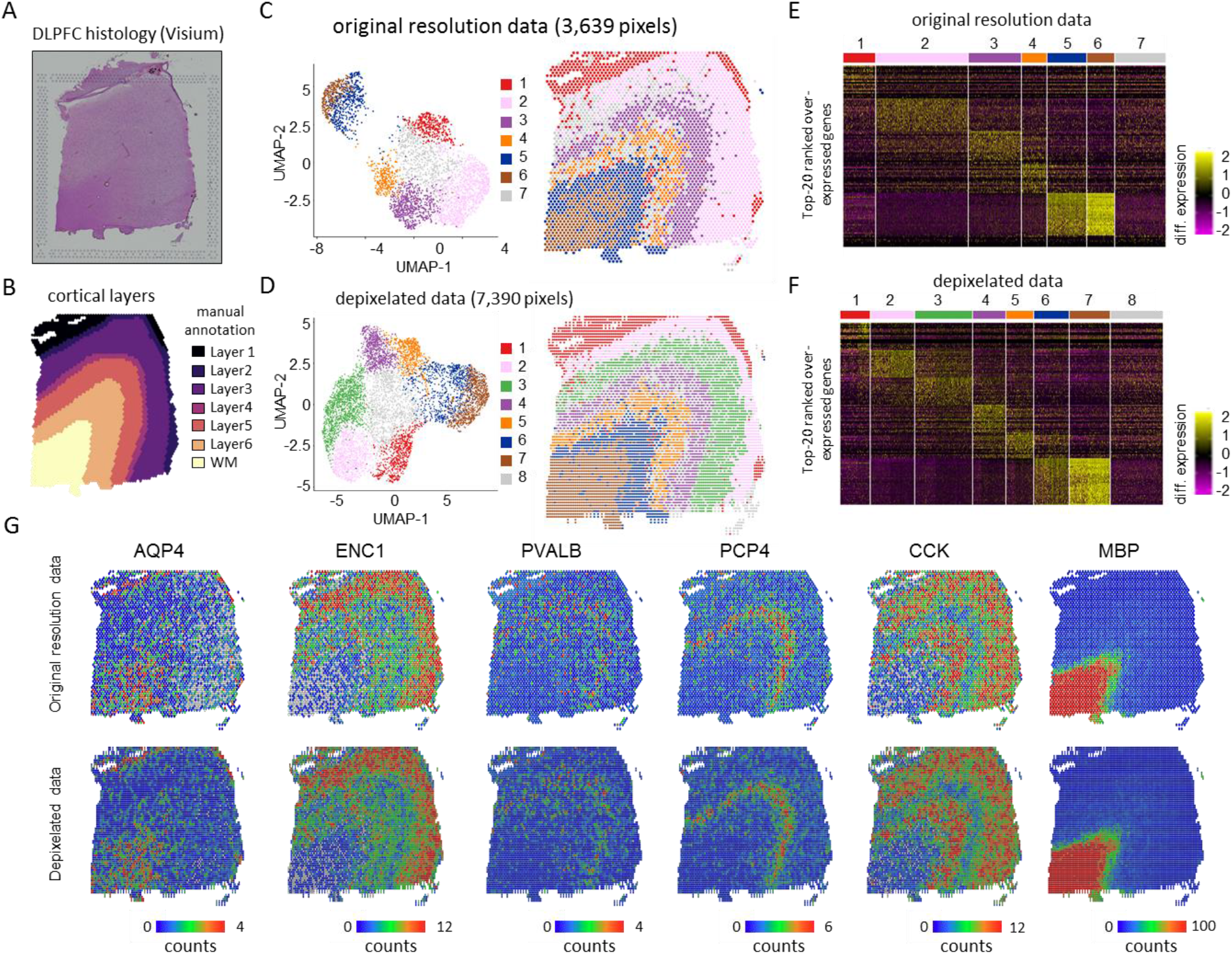
Depixelation data processing enhances the identification of known cortical layers in human brain datasets. **(A)** Visium slide presenting a dorsolateral prefrontal cortex (DLPFC) tissue section generated by (*Maynard et al*.,*2021*). **(B)** Manual annotation of the six cortical layers and the White matter (WM) to be retrieved in the Visium data, as illustrated in (*Zhao et al*.,*2021*). **(C)** Left panel: UMAP dimension reduction and clustering performed on the original Visium data composed by 3,639 pixels. Right panel: Projection of the seven identified clusters across the tissue map. **(D)** Similar to (C) but performed after a depixelation processing of the data leading to 2-fold gain in resolution (7,390 pixels). Notice that the depixelated data gave rise to 8 clusters, notably presenting the split of the cluster 2 of the original resolution data in two clusters (cluster 2&3 in depixelated data). **(E)** Heatmap displaying the top-20 ranked over-expressed genes per cluster retrieved in the original resolution data. **(F)** Similar to (E) but assessed on the depixelated data. **(G)** Six selected genes, known to be over-expressed at the different cortical layers, visualized on the grounds of their read counts retrieved either in the original resolution data, or after the depixelation processing.

A dimensionality reduction processing applied to the original resolution data (containing 3,369 pixels) followed by automated clustering allowed to identify seven clusters (**Figure 3C**). Similarly, after applying the spatial omics depixelation strategy, we obtained an enhanced spatial map composed by 7,390 pixels, which gave rise to 8 clusters after the dimensionality reduction process. A spatial projection of the identified clusters in the depixelated map revealed a clear stratification into 6 consecutive layers and the white matter region (brown cluster, **Figure 3D**). In contrary, projection into the tissue of the 7 clusters found in the original resolution data revealed a rather sparse view of 5 consecutive layers and the withe matter (**Figure 3C**). Notably cluster 2 (pink) in the original resolution map has been split in two layers (cluster 2 and 3) in the depixelated map (**Figure 3C&D**), in agreement with the manual annotation provided by the authors of the original study, but also with the data stratification obtained by BayesSpace (Zhao *et al*., 2021).

A heatmap analysis of the top-20 ranked over-expressed genes per cluster revealed that both cluster 7 in the original resolution data and cluster 8 in the depixelated map presented rather low gene expression levels, which explains their scatter behavior across the spatial maps, indicative for their irrelevant role for defining brain cortex layer subregions (**Figure 3E&F**). To further illustrate the tissue stratification performance, we have selected 6 genes described as being over-expressed in a layer-specific manner in the original study (Maynard *et al*., 2021). The gene coding for the myelin basic protein (MBP) appears specifically and strongly over-expressed in the withe matter layer, and it is well defined in both the original resolution map and the depixelated data (**Figure 3G**). The genes ENC1 (ectodermal-neural cortex 1) and CCK (Cholecystokinin), known to be associated to multiple layers in the brain cortex with exception to the with matter, presented a sharpen gene over-expression view in the depixelated data, revealing a main enrichment in layer 2 for ENC1 and layers 2 and layer 6 for CCK (**Figure 3G**). The gene coding for the Purkinje cell protein 4 (PCP4), known to be enriched in layer 5, appeared clearly enriched in both the original resolution and depixelated data, but the depixelation process appears to decrease the noisy enrichment signatures observed across other layers in the original data (**Figure 3G**). Finally, two genes presenting low expression levels in this data, PVALB and AQP4, presented an attenuated noise pattern behavior after depixelation process, with AQP4 being enriched in layer 1 and PVALB in layer 4; in agreement with previous data (Maynard *et al*., 2021).

### 3.3 Combining Low resolution data production with pseudo-interstitial pixels inference

The main advantage provided by unbiased strategies in spatial omics is the possibility to identify new gene players participating at the tissue architecture. This is possible thanks to the combination of next-generation sequencing (NGS) with molecular biology methods for capturing the target of interest across tissue sections. As consequence, these NGS-based approaches are systematically expensive because the discovery power is based on the afforded sequencing coverage, but also in the resolution of the spatial omics technology in use.

Previously we have described a cost-effective methodology for performing spatially-resolved transcriptomics assays by using in-house manufactured double-barcoded DNA arrays (Lozachmeur *et al*., 2023). This strategy relies in the manufacturing of DNA arrays by deposition of DNA probes covering a spot diameter of 100 microns, and a pitch distance of 177 µm (**Figure 4A**). While presenting a lower resolution than the current commercially available solutions, this technology allowed us to perform – in a cost-effective manner - several consecutive spatial transcriptomics assays across human brain organoids, giving us the possibility to generate a 3-dimensional view of the various genes getting over-expressed across the tissue (**Figure 4B&C**).

**Figure 4.**
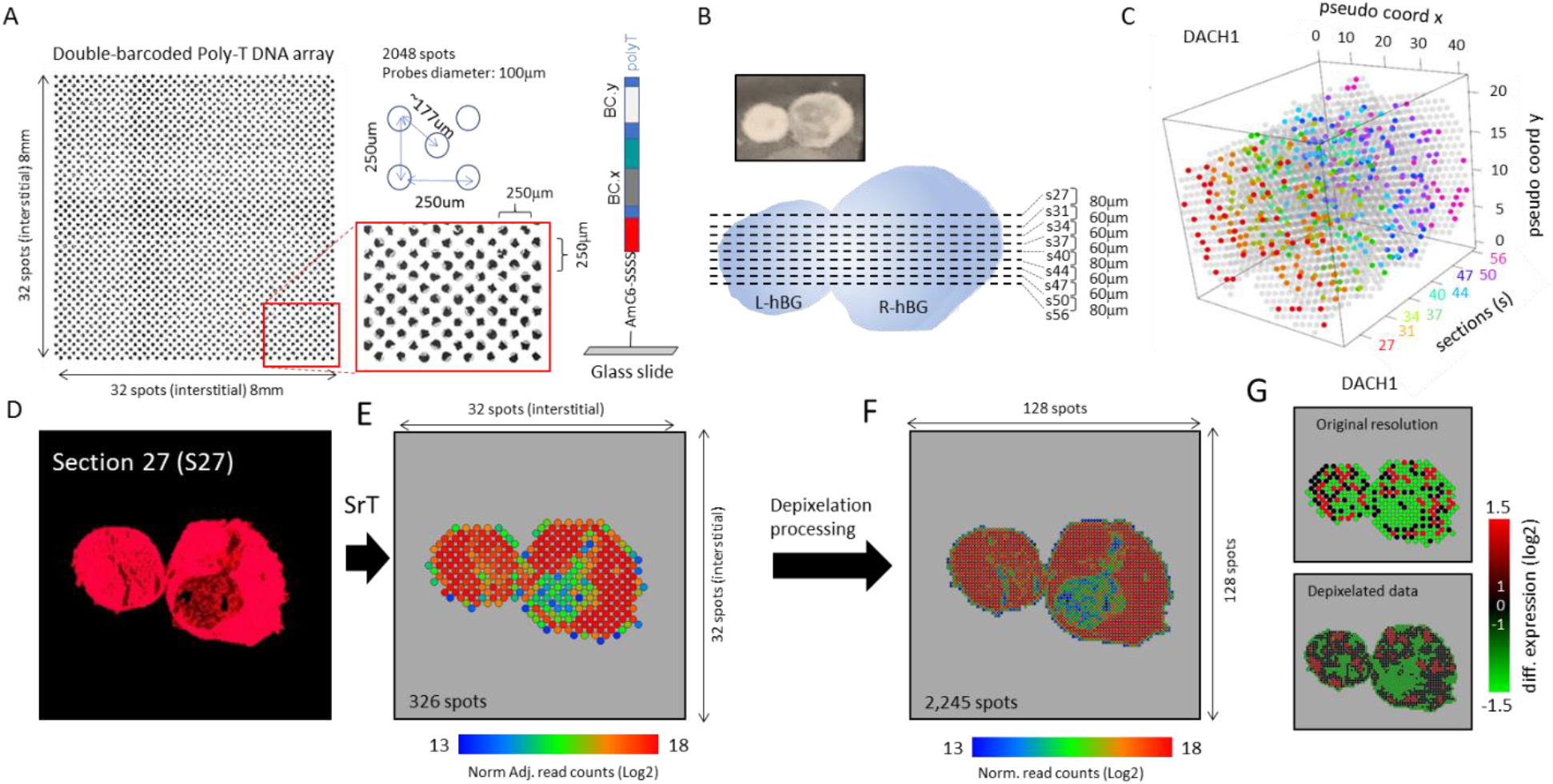
Combining Low resolution data production with pseu-do-interstitial pixels inference. **(A)** Micrograph displaying a DNA array manufactured in-house presenting two molecular barcodes; a molecular barcode defining the X coordinate (BC.X) and a second barcode defining the Y coordinate (BC.Y) of the printed probe. Manufactured DNA arrays present a 100 microns spot diameter, and a pitch distance of 177 µm. The displayed DNA array present poly-T sequences at the end of the probes for capturing messanger RNA. **(B)** Top: Micrograph displaying two human brain organoids (hBG) cryo-sectioned together for spatial transcriptomics assays. Bottom: Scheme illustrating the collection of 9 consecutive sections across both hBGs (s27-s56). L-hBG & R-hBG: Left and Right human brain organoids. **(C)** Three-dimensional view displaying the over-expression detection of the gene DACH1 (Dachshund Family Transcription Factor 1) in the different spatially-resolved transcriptome maps assessed from the collected sections. **(D)** Micrograph displaying the section 27 where the produced cDNA during the spatial transcriptomics preparation has been labelled by dCTP-Cy3 incorporation. **(E)** Spatially-resolved Transcriptomics (SrT) map composed by 326 pixels or spots. SrT map has been first quantile normalized for correcting for potential technical variations, then adjusted for tissue signal intensity, retrieved in (D), for taking in account difference in cell density. **(F)** SrT map after applying the depixelation process (3 round of expansions), leading to a total of 2,245 pixels or spots; i.e. a ∼7-fold gain in resolution. **(G)** DACH1 over-expression levels displayed either in the original resolution data or after the depixelation process.

As illustrated in **Figure 4D**, the reprocessing of this published data by incorporating a step of data depixelation (3 expansion steps) allowed to convert a spatial map composed of 326 spots (or pixels) into a denser image presenting 2,245 pixels, thus representing a ∼7-fold gain in resolution (**Figure 4D-F**). As consequence, the obtained depixelated map allows to better resolve spatial regions presenting local gene over-expression signatures, as illustrated in the case of the gene DACH1 (Dachshund Family Transcription Factor 1); known to be highly expressed in the proliferating neuroprogenitors during human neocortical and striatal development (Castiglioni *et al*., 2019), and recently detected in other brain organoids progression studies (Chiaradia *et al*., 2023) (**Figure 4G**).

Overall, by combining a cost-effective spatial omics technology with a data depixelation processing, we can obtain high resolutive information presenting the required gene discovery power issued from the use of NGS-based technologies.

## 4. Conclusion

In this study we describe Multilayer-expander, a computational strategy allowing to gain several folds of data resolution by inferring pseudo-interstitial pixels from the information available in their nearest neighbors. This strategy is fully inspired by current methods used form image-depixelation processing applied in large number of applications. Indeed, spatial omics data can be considered as a color image, with the main difference that pixels in classical color images are composed by three values (R,G,B); while spatial omics data are composed by Gexels (Gene expression elements) composed by several thousand of gene expression read counts. Hence, exploiting the large know-how used in the field of image processing can become beneficial for the field of spatial omics. In this context, we consider Multilayer-expander as a pioneer tool using the concepts behind the up/down-scaling of images for different screen resolutions, notably by Nearest Neighbor resampling (reviewed in (S Malini and Patil, 2018)), anticipating future implementations focused in more developed strategies.

## 5. Acknowledgements

We thank all members of the team SysFate for contributing to the discussion of this project.

## 6. Funding

This work was supported by the institutional bodies CEA, CNRS, Université d’Evry-Val d’Essonne. A. G and M.D are supported by EU grant HORIZON N° 101070740.

### Conflict of Interest

none declared.

## Notes

### Competing Interest Statement

The authors have declared no competing interest.

